# Meggie – easy-to-use graphical user interface for M/EEG analysis based on MNE-python

**DOI:** 10.1101/2022.09.12.507592

**Authors:** Erkka Heinilä, Tiina Parviainen

## Abstract

**Background:** In the last decades, electrophysiological imaging methodology has seen many advances and the computational power in the neuroscience laboratories has steadily increased. Still, the new methodologies are unavailable for many. There is a need for more versatile analysis approaches for neuroscience specialists without a programming background.

**New method:** Using a software which provides standard pipelines, provides good default values for parameters, has a good multi-subject support, and stores the used analysis steps with the parameters in one place for reporting, is efficient and fast. In addition to enabling analysis for people without background in programming, it enables analysis for people with background in programming but a limited background in neuroscience. When constructed with care, the GUI may guide the researcher to apply analysis steps in correct order with reasonable default parameters.

**Comparison to existing methods:** Two existing software, EEGLAB and Brainstorm, both provide an easy-to-use graphical user interface for end-to-end analysis for multiple subjects. The key difference to work presented here is the choice of language. The scientific community is moving *en masse* towards the python programming language, thus making it an ideal platform for extendable software. Another problem with Matlab is that it is not free - both from the perspective of open source and concrete monetary resources. Within the current trend towards increasing open science, covering data, analysis and reporting, the need for open source software is imperative.

**Results:** Meggie is an open source software for running MEG and EEG analysis with easy-to-use graphical user interface. It is written in Python 3, runs on Linux, macOS and Windows, and uses the MNE-python library under the hood to do heavy lifting. It is designed to allow end-to-end analysis of MEG and EEG datasets from multiple subjects with common sensor-level analysis steps such as preprocessing, epoching and averaging, spectral analysis and time-frequency analysis. Most of the analysis steps can be run for all the subjects in one go, and combining the results across subjects is made possible with grand averages. We have emphasized the extendibility of Meggie by implementing most of the Meggie itself as plugins, thus ensuring that new plugins have access to all necessary core features.

**Conclusion:** Meggie answers the demand for easy-to-use and extendable python-based graphical user interface that provides an end-to-end analysis environment for M/EEG data analysis. It is freely available at https://github.com/cibr-jyu/meggie under the BSD license. Installation instructions, documentation and tutorials are found on that website.

**Highlights:**

- MNE-python-based graphical user interface for M/EEG analysis
- Easy to use even without programming background
- Has good support for multiple subjects
- Written in python, and can be easily extended with plugins
- Free and open source with BSD license

## Introduction

The first electroencephalography (EEG) was measured almost 100 years ago in 1924 by Hans Berger. The stereotyped responses to stimuli, i.e. event-related potentials (ERPs), were discovered in 1930s, but truly became a popular method for investigating brain activity in the 1960s when the computerized solutions, such as CAT (the computer for average transients), for averaging over multiple responses became available. ERPs were simple enough to be computed in the advent of computer science and yet were useful in the study of the human brain (Morgan-Short and Tanner, 2014).

Since then, electrophysiological imaging methodology has seen many advances. The arrays of electrodes in a single Electroencephalography (EEG) cap has grown steadily in size (and are, nowadays, often in hundreds) increasing the spatial accuracy and the signal quality.

Magnetoencephalography (MEG), the recording of magnetic signals originating from the electrical activity in the brain, has been around since the 1980s and even though more expensive than EEG, is becoming more and more popular for its clean signals and good spatial accuracy (Hari and Salmelin, 2012).

In the neuroscience laboratories, the computational power has steadily increased. Methods developed within electrical engineering such as time-frequency analysis and data-driven characterizations of the signal content have been introduced to M/EEG analysis as well. Not every aspect of the brain can be captured with only ERPs, as there can be activity in the brain that is not strictly time-locked to the stimuli, or emerge even without external stimulation. Indeed, brain signal processing is essentially characterized by both exogenous (i.e. responses evoked by external stimuli) and endogeneous (i.e. activity emerging due to internal computation not necessitating external triggers). It is only in the 2000s when the tools to do more than just ERP analysis have become widely available (Li et al., 2013; Morales and Bowers, 2022)

Currently, the state-of-the-art methodology is developing rapidly. In addition to ERPs, resting state analysis based on spectral decompositions and induced response analysis with wavelets are considered standard. More accurate localization of the signals via inverse modeling is increasingly popular and machine learning ideas, such as decoding and deep learning, are being introduced to neuroscience (Zhigalov et al., 2019). At the same time, even the standard methods are not always easily available for a researcher. In the current software landscape there are two popular tool boxes for M/EEG analysis. Fieldtrip provides an extensive list of analysis methods for a user able to write matlab scripts and MNE-python enables a large repertoire of analysis methods for python users (Oostenveld et al., 2010). However, the learning curve in mastering these command-line-based environments is quite strongly dependent on the background experience of the researcher. While academics with computational background can quickly adopt M/EEG specific use of a programming language, some other background training, e.g. psychology, linguistics, or neurobiology, does not equip students with needed knowledge to design and choose the best pipelines for neuroscientific data analysis. For a more efficient use of researcher’s time, there should be tools that can be tweaked and adjusted without having to learn the inside secrets of a programming language. Moreover, even the computer scientist may still prefer to use an efficient graphical user interface (GUI) sometimes, as besides providing an easy interface to the computations, they can also be designed to guide the correct progression of analysis, with variable level of freedom in selecting specific parameters at each stage.

The problem has been approached before. EEGLAB (Delorme and Makeig, 2004) is a widely used software, based on matlab language, that is used mainly for EEG data with a graphical user interface. Matlab-based Brainstorm (Tadel et al., 2011) software, on the other hand, provides graphical user interface for mainly MEG, but also EEG analysis. There is also a new mne-python based graphical user interface, mnelab, providing access to mne-python functionalities in a programmatic manner, i.e explicitly wrapping each of the mne functions in the user interface. Although the existing GUI’s for electrophysiological brain imaging data analysis already bring significant impact for the needs of the fields, some problems are still worth addressing.

The scientific community is moving *en masse* towards the python programming language. Python is the language of choice for machine learning applications, and thus the software, that is to provide access to that vault also for neuroscientists, should be implemented in python. Another problem with matlab is that it is not free - both from the perspective of open source and concrete monetary resources. Within the current trend towards increasing open science, covering data, analysis and reporting, this can cause increasing challenges for neuroscientists. While python-based mnelab is a well designed software and faithful to mne workflows, it does not profile as a Brainstorm-like all-encompassing end-to-end software. That is, it tries not to provide an analysis environment where you can add all your raw datasets, run sequential analysis steps on them in a parallel manner, and then combine them to get the final quantity of interest.

Meggie is an open source software for running MEG and EEG analysis with easy-to-use graphical user interface. It is written in Python 3, runs on Linux, macOS and Windows, and uses the MNE-python library (Gramfort et al., 2014) under the hood to do heavy lifting. The project started in Jyväskylä Centre for Interdisciplinary Brain Research (CIBR) in 2013 and has been used in-house since the early days. It became publicly available in 2019 and was officially released in 2021. Meggie was developed for the cognitive neuroscience research community, consisting of students and academics with variable background training and level of experience in working with programming languages, to be able to apply the state-of-the art MEG analysis protocols both in experimental work and in education.

There are two key design ideas behind Meggie. First, in order to respond to the need for an increased number of subjects in neuroimaging, experiments with multiple datasets (subjects) need to be well supported. Thus, Meggie allows adding all the data of all subjects of an M/EEG experiment to the software, and as you run the analysis steps, you can almost always run them for all subjects in one go. Second, to provide guiding principles for the analysis pipeline also for beginning users, the functionalities in Meggie are organized in analysis pipelines so that it is easy and clear to run analysis steps sequentially, without getting lost in the way. For example, in the ERP analysis, the data is imported and preprocessed, the epochs are created and averaged, and the resulting time courses are visualized and compared across different conditions. The results can be saved either in image or text (numeric) format for further statistical analysis. In the meggie user interface, this corresponds to moving through a sequence of tabs, or from top to bottom within a tab, making it intuitive to understand the progression of the data analysis steps. The pipelines have been designed to comply with the recent guidelines for M/EEG analysis (Gross et al., 2013). To combine these two needs posed by the analysis requirements of large-scale M/EEG samples, the underlying principle is to guide the user to process the data in ‘waves’ from raw signals, through the analysis steps, to the final results.

Thus, Meggie is designed to allow end-to-end sensor-level analysis with common analysis steps such as preprocessing, spectral analysis of continuous data segments, epoching and averaging, and time-frequency analysis. Most of the MEG and EEG formats are supported in the input phase and the processed data can be saved either as csv files or in native formats (e.g fif).

Meggie implements an extensive plugin system, and except the very core, most of the Meggie itself is implemented as internal plugins. Plugins allow extending the functionality via python packages external to Meggie itself, and can be developed and used in-house or shared with others. Some plugins, for e.g source analysis and statistical analysis via cluster permutation tests, can already be found from the software website.

## Methods and results

### Overview

The main view of Meggie consists of the left column containing the *experiment* (Fig 2A) and *subject* (Fig 2B) information and the related functionalities such as setting channel groups or adding subjects to the current experiment, and the right column containing all the possible analysis *actions* organized in tabs (Fig 2C). There is a console at the bottom of the screen (Fig 2D), containing information about the current analysis session. The menu at the top can be used to create and open experiments, set preferences or examine previously applied actions. The Meggie-specific nomenclature is explained in Table 1.

**Table 1.** Summary of terms used in the software.

|  |  |
| --- | --- |
| Experiment | The main data structure of Meggie, which contains all the subjects. |
| Subject | The main container for raw data, and subject-specific results. |
| Datatype | The subject-specific intermediate results have a corresponding datatype. Such results are automatically stored in the experiment folder. By default, Meggie includes epochs, evoked, spectrum and TFR datatypes. |
| Experiment folder | A folder in the file system where experiment, its subjects, and all the subject-specific results are stored. |
| Pipeline | Pipeline is a set of actions relevant to some specific type of analysis. By default, Meggie contains pipelines for continuous data analysis, ERP analysis and induced response analysis (see Fig 1). |
| Action | A M/EEG analysis from start to beginning usually consists of many analysis steps, e.g. artifact removal, filtering, resampling, epoching and so on. Actions wrap these steps, and, in short, consist of a python function and relevant parameters. Use of actions is automatically logged and actions previously used within the experiment can be inspected from past actions dialog. |
| Plugin | A plugin is a python package that can introduce additional datatypes, actions and pipelines. |

The standard usage of Meggie starts by creating an experiment. At the creation, name and author, as well as a *pipeline*, are chosen for the experiment. Normally, only one pipeline is relevant for a specific research question, and the use of pipelines make it possible to simplify the UI for the specific task at hand, guiding the user to take correct steps (see Fig 1 for a schematic diagram of the pipelines). As a result, an *experiment folder* is created to the file system. It will store all the files and information related to the experiment. After this, subjects (i.e. M/EEG recordings of individual research subjects) can be added to the experiment using the “Add new..” button in the left column. From here onwards the actions provided in the tabs of the right column can be used to sculpt the datasets into the results. Most of the actions, such as filtering or creating epochs, can be applied to all subjects in one go.

**Figure 1.**
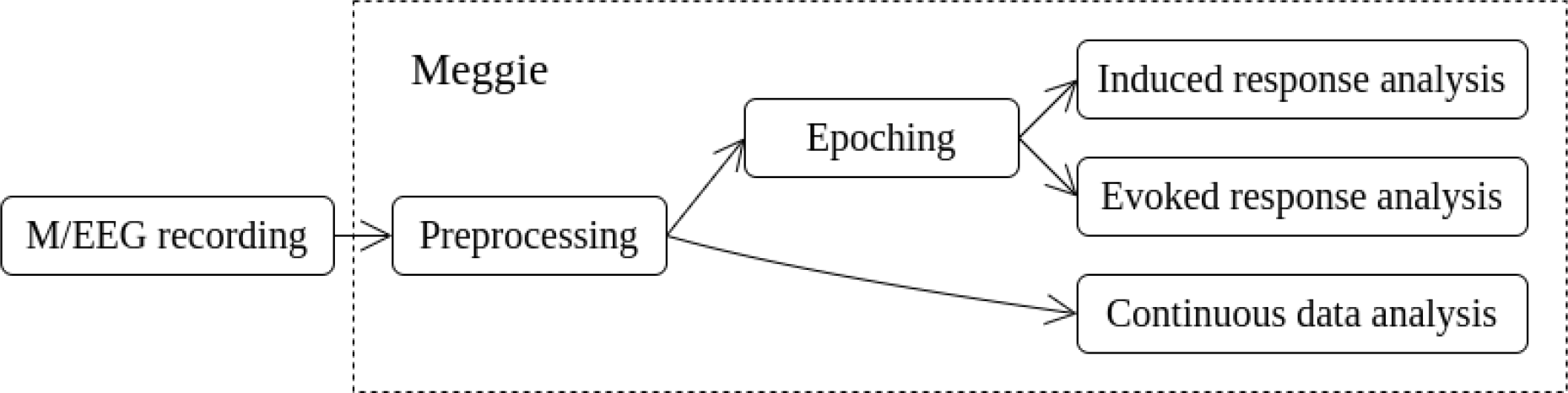
The three main pipelines in Meggie.

**Figure 2.**
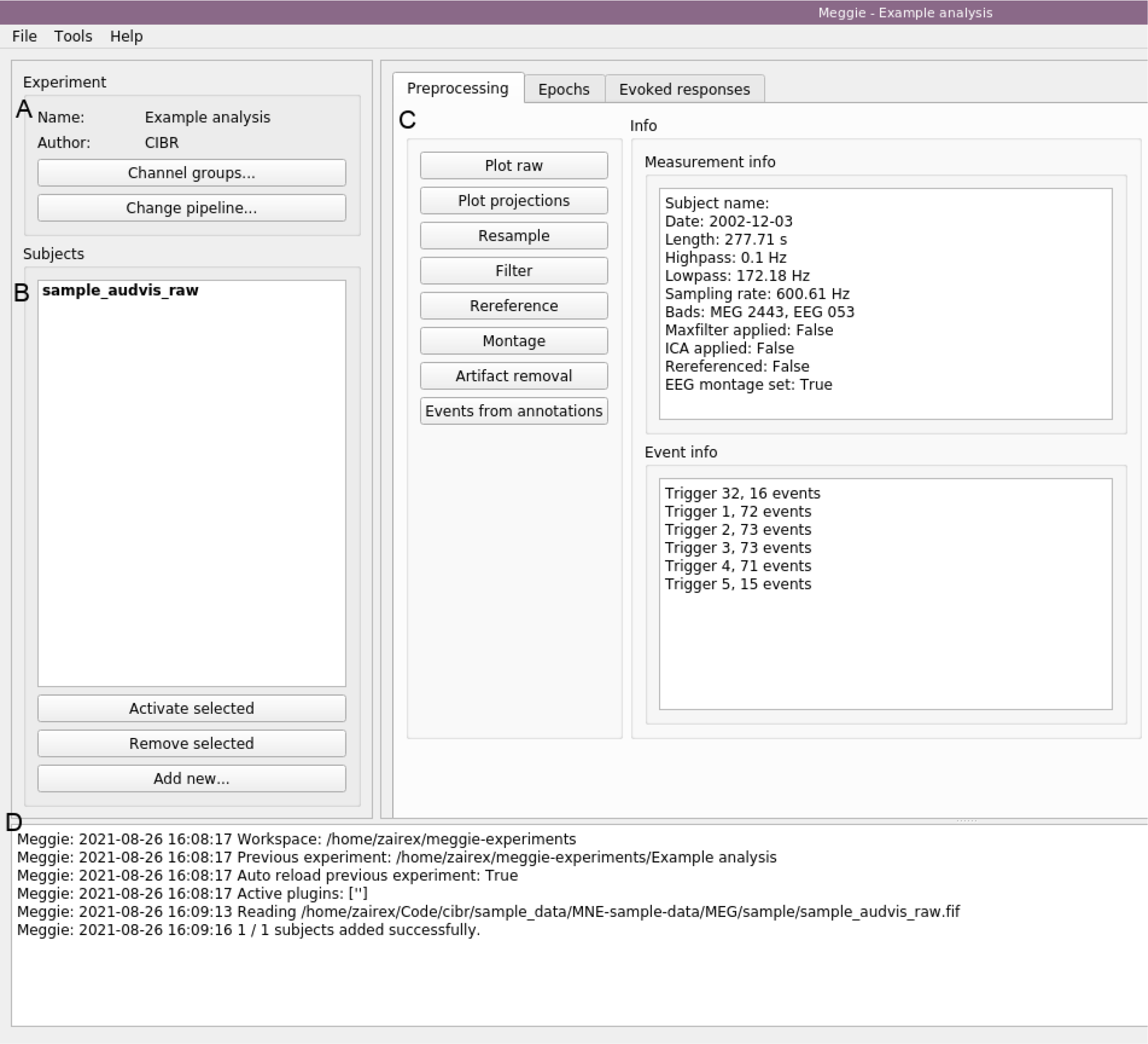
The main view of Meggie, with experiment section (A), subject section (B), analysis section (C) and console (D).

### Data import

Meggie uses mne-python’s functions to read M/EEG raw recordings. This means that at least the following formats should be supported: edf (.edf), bdf (.bdf), gdf (.gdf), brainvision (.vhdr), fif (.fif), eeglab (.set), cnt (.cnt), egi (.mff), eximia (.nxe), nirx (.hdr), fieldtrip (.at), artmemis123, (.bin), nicolet (.data), kit (.sqd) and ctf (.ds). As meggie is actively maintained, any new formats added to mne will be available in Meggie too. Internally the fif format is the format of choice in mne, and thus any input recording will be converted to that format.

Behind the scenes, when a subject is added to Meggie, the specified raw data is copied (not moved) to the experiment folder. This way, no harm can happen to the original files in the file system, as the subsequent actions that are run on the subjects will be based on the new files.

### Preprocessing

As an important feature at preprocessing level, Meggie supports simple data inspection through a time series plot. In addition to just looking at the raw data, the plot allows annotating data segments and setting bad channels.

In Meggie, artifact removal (i.e. removing noise or confounders such as eye blinks from the data) can be accomplished with independent component analysis (see Figs 3 and 4). ICA is a popular blind source separation method which transforms the channel data to a more interesting space where the signals, i.e. the components, are statistically independent from each other. For electrophysiological signals the most typical and robust components in M/EEG usually associate with eye blinks, and also the cardiac signal represents easily a discernible component in the ICA, especially in MEG. After the transformation, the resulting components can be inspected (see Fig 4), and the artifacts (i.e the blinks and cardiac signals) can be selected for removal (see Fig 3A). Before subtracting the artifacts from the original signals, it is possible to inspect the difference between the original and the artifact-free signals (see Fig 3B).

**Figure 3.**
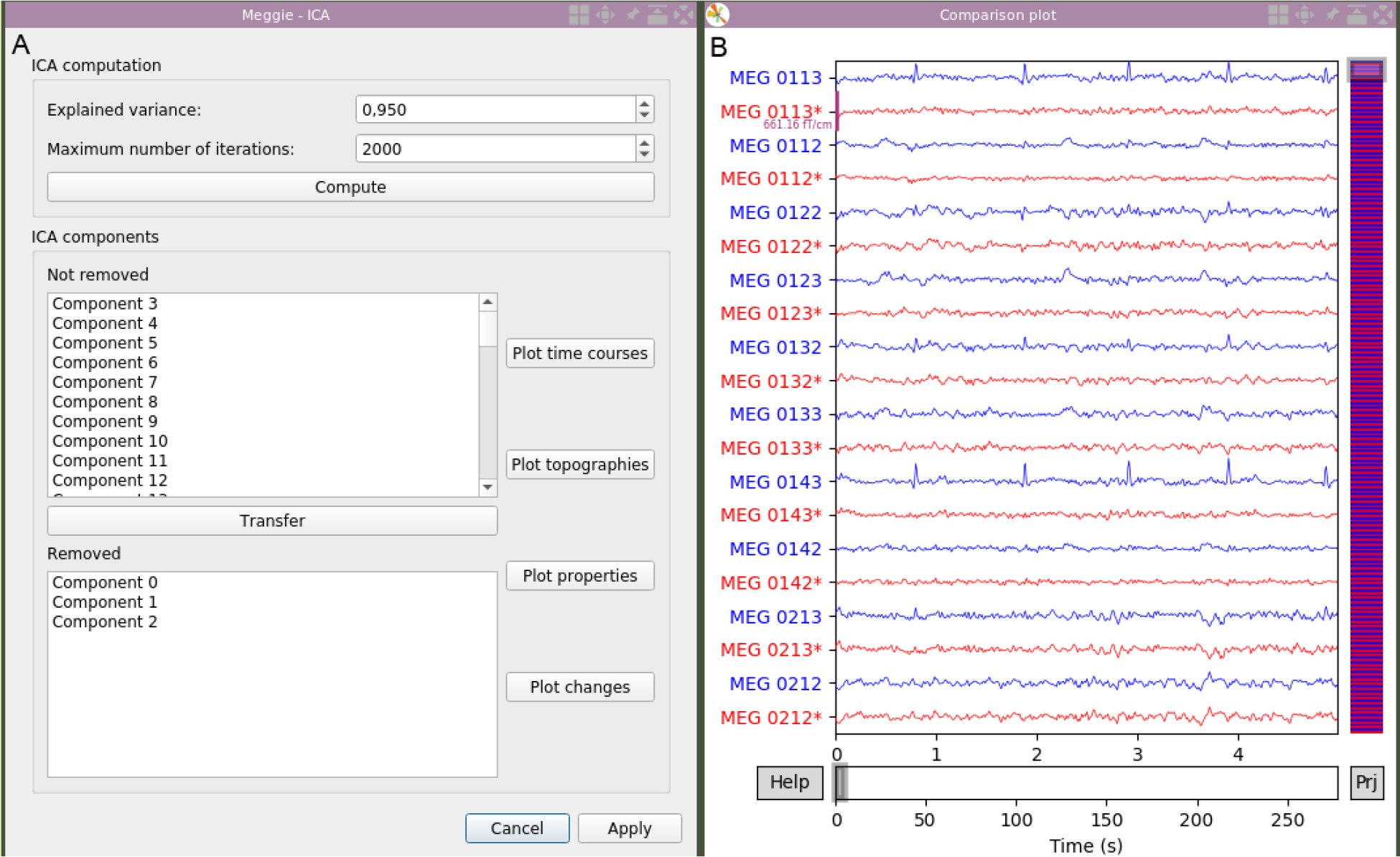
Artifact removal dialog (A) and a comparison plot (B), which is opened with “Plot changes” in A.

**Figure 4.**
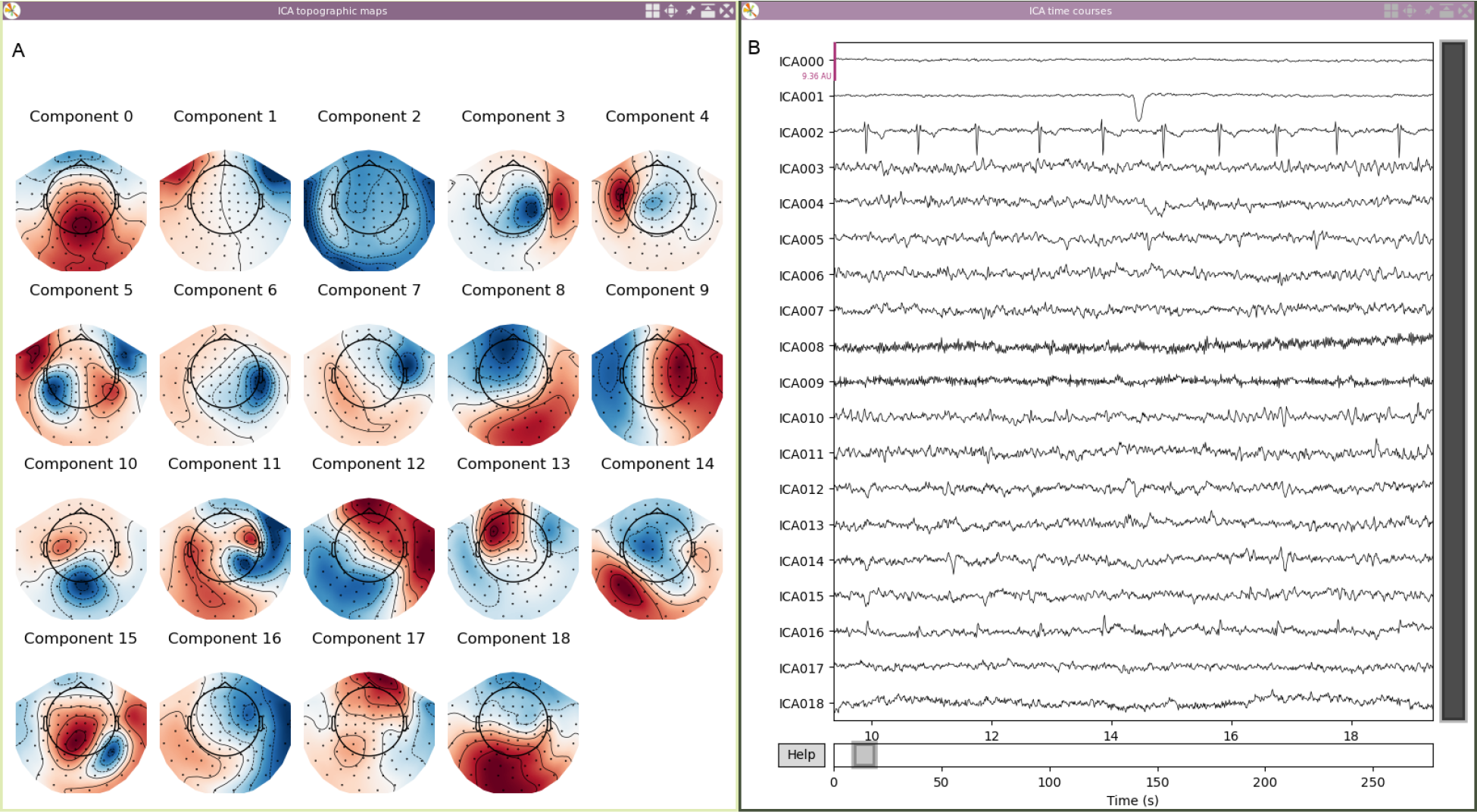
Topomap (A) and time series (B) plots of independent components.

Another way to improve the data quality is with frequency filters. Applying a frequency filter to attenuate some wavelengths, where the signal of interest should not be present, increases the signal-to-noise ratio of the remaining signal. Three basic FIR-filters are available in Meggie: a low-pass, a high-pass and a notch filter (i.e. band-stop). Using a high-pass filter even with a cut-off as low as 0.1Hz, can improve the stability of ICA, which is sensitive to slow drifts in the signal. A notch filter can be used to remove certain specific narrow frequency bands such as the line noise.

To limit the computational time in computationally demanding analysis steps, such as wavelet transformations, or to limit the storage requirements, it may be necessary to downsample the data. Downsampling by a factor of N means picking every Nth sample. To avoid aliasing effects, the data is automatically low-pass filtered to half of the new sampling rate. Note that this can somewhat distort the event timing, and should be used with care if high temporal accuracy is needed.

Meggie thrives to support both the EEG and the MEG data in a seamless uniform way. However, even though MEG recordings are embedded with the sensor locations, this is usually not the case with EEG data. Thus, to facilitate plotting of topographic maps for EEG data that does already contain sensor locations, Meggie provides a way to add a default montage (i.e. locations of the sensors in the 3d space) to the dataset either from a list of standard montages (such as Biosemi or EGI) or from a file. Also a way to change the referencing of the EEG channels, either to a certain channel or to an average, is provided.

Internally, Meggie prefers events over annotations. Events are points in time that also have an integer value signifying what happens at that point. Annotations are segments in time that have a starting point, an ending point and an associated value in text. As there are legitimate use cases for annotations and the raw data plot supports annotation of data, we provide an utility to convert annotations to events. The available preprocessing actions are also visible in Fig 2C.

### Pipelines and visualization

The goal of analysis is often to transform the data from raw temporal signals to signals or numeric values that summarize the essential features of the data given the research questions. In Meggie, three basic pipelines are provided: continuous data analysis through spectrums, evoked response analysis through epoching and averaging, and induced response analysis through time-frequency representations (TFRs). All of the pipelines begin with the preprocessing stage and then continue to actions specific to them (see Fig 1 for a schematic diagram).

### Evoked responses

Because of the sensor noise and the complex nature of the brain activity, time-locked brain responses are investigated by collecting many samples (epochs) of the brain response to a specific stimulus, and then averaging them together. This way, the activation not related to the stimulus should vanish, as it is expected to have a mean of zero, and the averaging decreases variance. However, the activation time-locked to the stimulus should not vanish, as it is assumed to have a non-zero mean. This increases the chance to see a pattern of brain activation otherwise hidden in the other task-unrelated activity or noise (Thigpen and Keil, 2017).

To do this in meggie, you first create an epoch collection in the Epochs tab. These are based on event information that should be embedded into raw data. A versatile utility for event selection is provided with the possibility to, for example, do event code masking and add delays (see Fig 5A). The resulting collection can be inspected. See Fig 5B for one kind of visualization. Then, in the Evoked responses tab, it is possible to average the collection of epochs together. Here, the responses can be investigated with many visualization methods, such as channel-wise plots, channel average plots and topographic maps. See Fig 6 for an example of an overall auditory response in a single subject visualized as a topographic map evolution for the three channel types present in the data.

**Figure 5.**
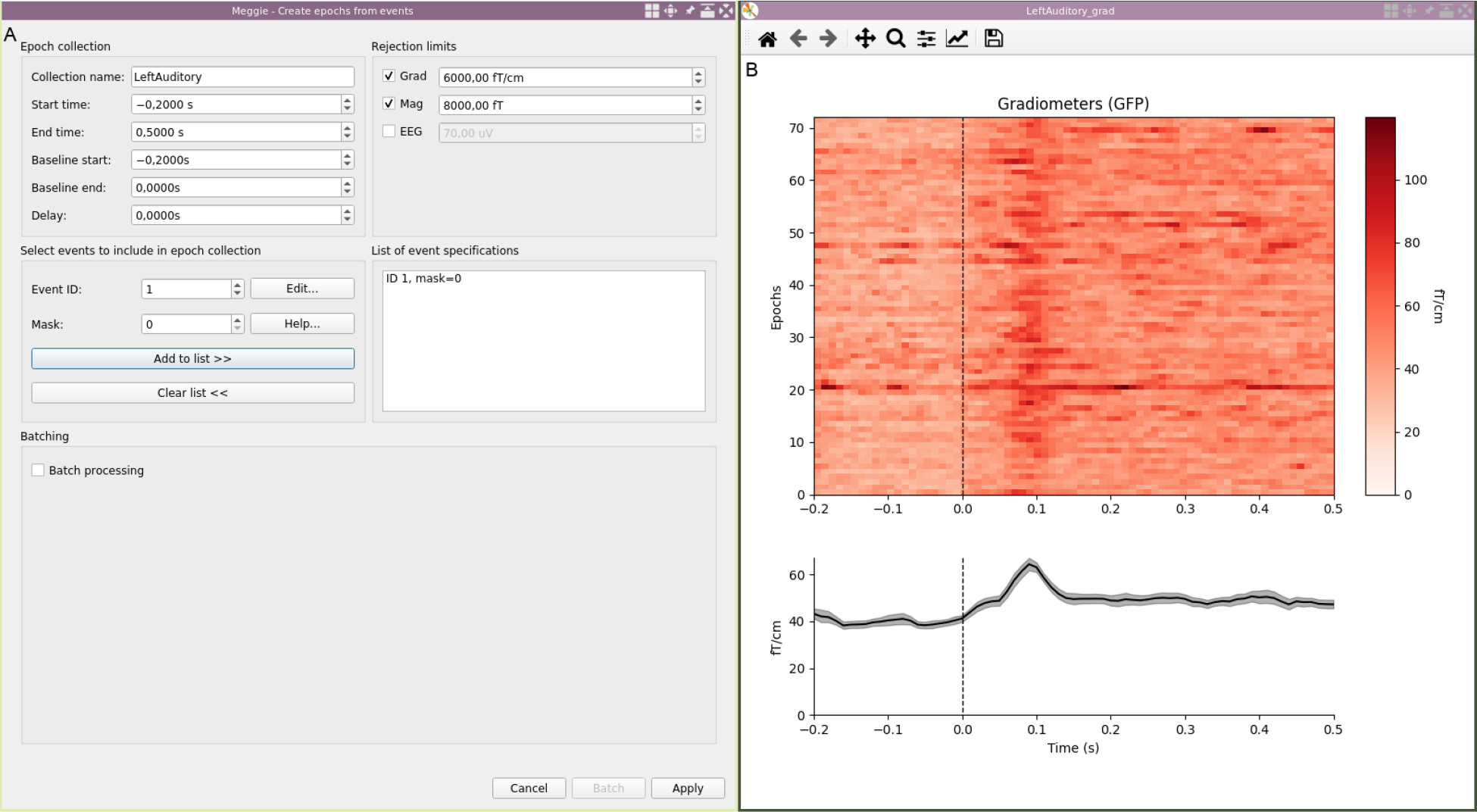
Epoch creation dialog (A) and an epochs image plot (B).

**Figure 6.**
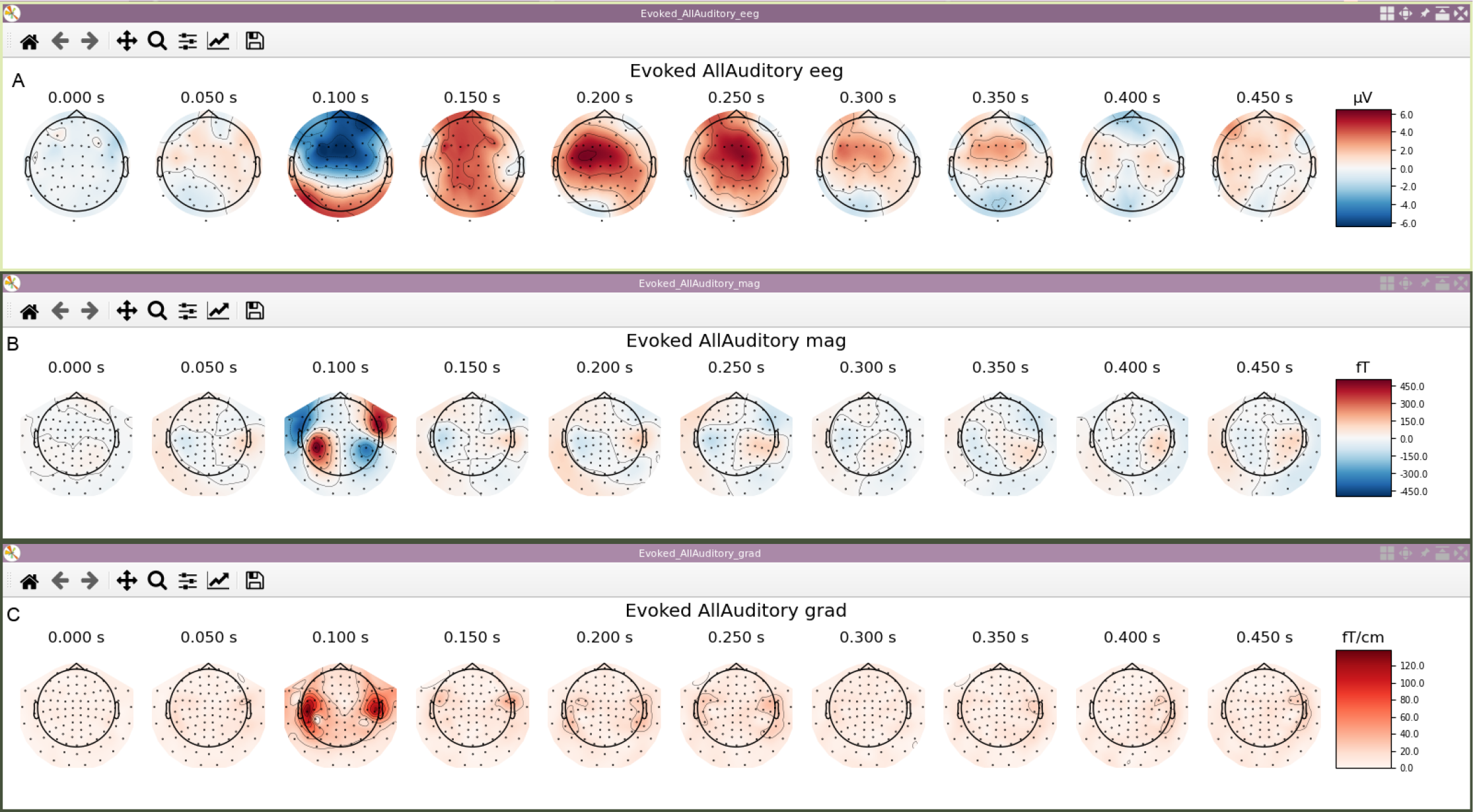
Topographical maps of evoked responses for EEG channels (A), magnetometers (B) and gradiometers (C) visualized over time.

### Induced responses

Sometimes, we are interested in brain responses that are related but not strictly time locked to (and hence ‘induced’ by) the stimulus. In this case, averaging may make the signal of interest vanish too. One example of such a signal is a brain oscillation. If the amplitude of oscillation is modulated by the stimulus, but the phase is kept intact, the raw signal may vanish in averaging, but have a non-zero spectral amplitude. In this case, it is possible to first compute a dynamic representation for the power content for each epoch in the collection separately, and then average these representations together to increase the signal-to-noise ratio as before. In Meggie, wavelets are used to extract a time course for each of the frequencies of interest separately, which are then stacked together to create a time-frequency representation (Morales and Bowers, 2022).

Time-frequency representations can be visualized as heatmaps, as shown in Fig 7. In the figure, we also demonstrate the use of custom channel groups to average TFRs, computed for each channel separately, over a specific group of channels. The figure shows response to somatosensory stimulus for “occipital” and “vertex” channel selections. As the comparison of two or more TFRs (for example, for three different stimuli) can be difficult (as they are difficult to plot over top of each other), Meggie provides a simple way to compare TFRs for a specific narrow frequency band. In this case, the frequencies of interest can be averaged together and only a time course remains. These are called temporal spectral evolutions (TSEs) and can be drawn within the same plot for easy comparison (Vanni et al., 1997).

**Figure 7.**
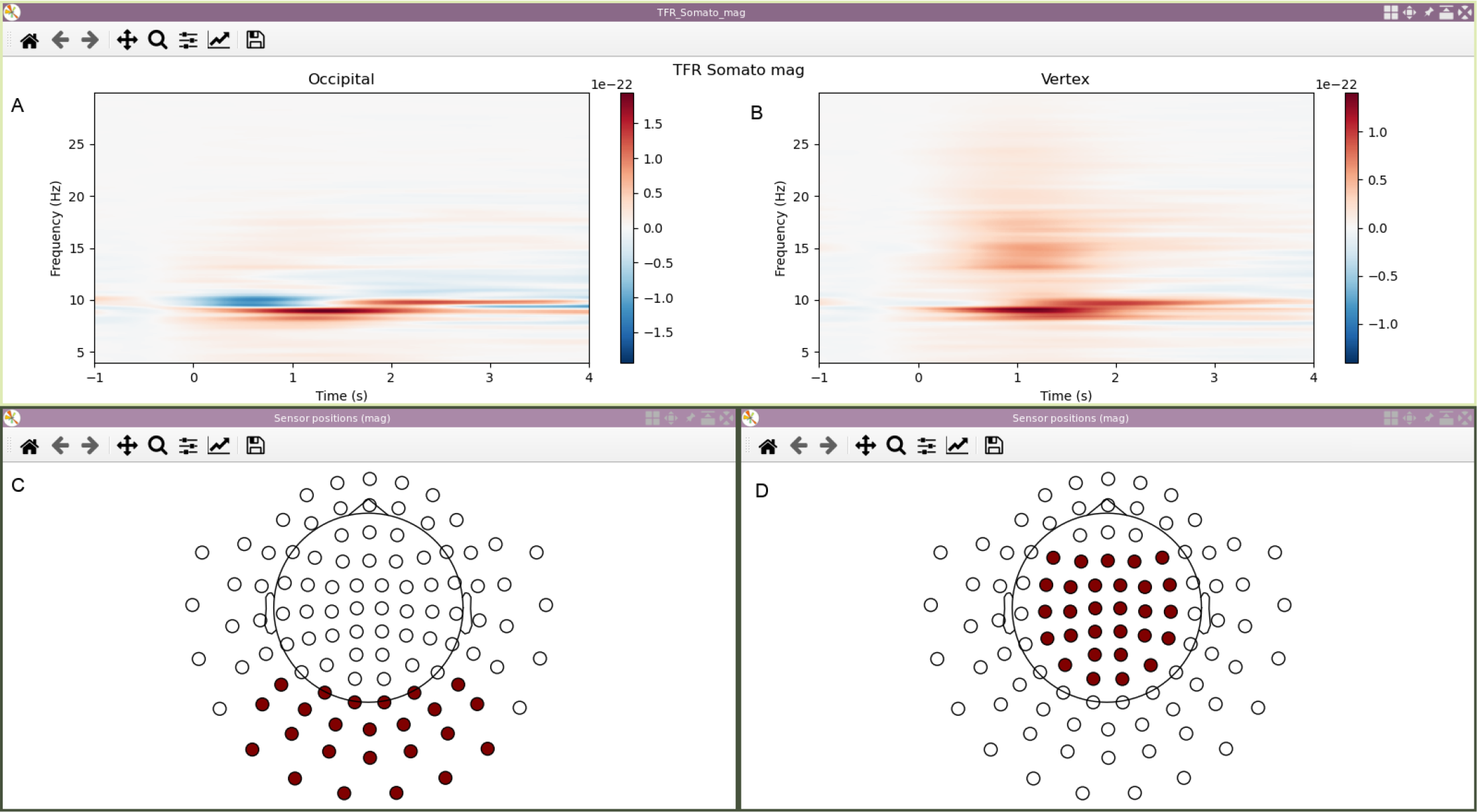
TFR’s averaged over custom channel groups. Plots on the left show TFR of the somatosensory response (A) averaged over occipital channels (C). Plots on the right show TFR of the somatosensory response (B) averaged over vertex channels (D).

### Analysis of continuous signals

If data is of continuous nature, i.e there is no stimuli to base the epochs on, the previous methods can not work. However, it is still possible to analyze the average spectral content of the continuous signals. This means decomposing the signals to their frequency components (analogous to light which consists of different colors). Such ‘spectrums’ are usually computed with Welch’s method, which utilizes a series of Fourier Transforms under hood (Dressler et al., 2004; Hohaia et al., 2022; Zhao and He, 2013).

Meggie provides a versatile utility for selecting time intervals of interest. They can be based relative to event information or relative to the start or the end of the measurement. As for evoked responses and induced responses, spectrums can be visualized channel-wise or as channel averages. Conditions of interest can be visually compared, and saved for further analysis. See Figure 8B for two spectrums visualized in a single plot. The red curve is based on eyes closed resting state while the blue curve is based on eyes open resting state. The spectrums were averaged over occipital channels (as in Fig 7C) and over 28 different subjects. Figure 8A shows a csv file containing data from all 28 subjects separately opened in a datasheet program.

**Figure 8.**
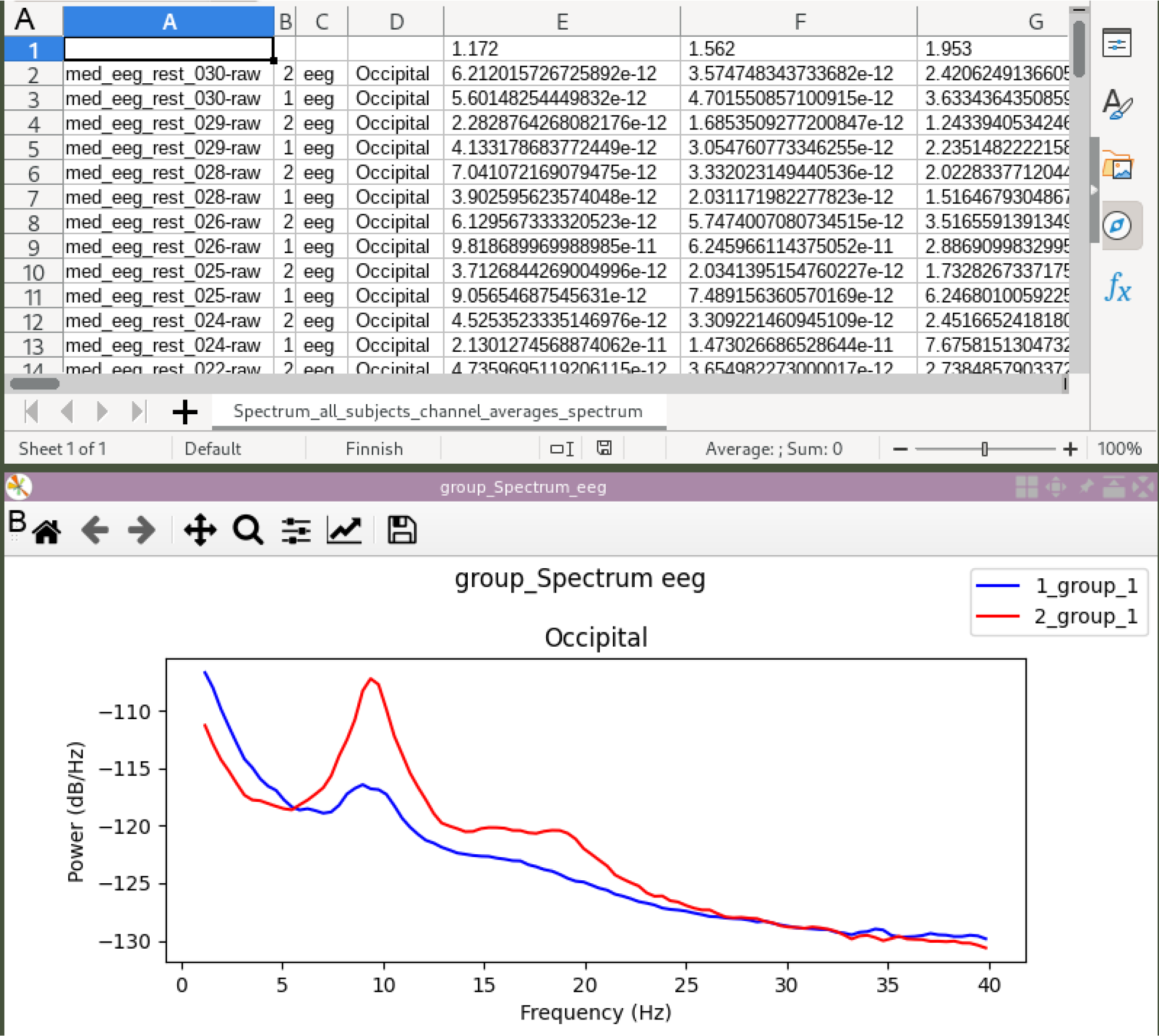
A datasheet saved with Meggie (A) and a matching occipital channel average plot of eyes-open (blue curve) and eyes-closed (red curve) resting state conditions averaged over 28 subjects (B).

### Data output

The figures created in Meggie are matplotlib figures, and can be tweaked with the graphical user interface of matplotlib library. If more control over figures is needed, the data can be saved as structured text (csv) files, and visualized in external software such as Excel or Libreoffice. Exporting to csv also makes it possible to, for example, use external software such as SPSS for statistical analysis. Figure 8A shows spectral data saved in Meggie and opened in Libreoffice. Both the figures and the csv files can be saved either for all the available channels or, if applicable, the data can be averaged over groups of channels, and the channel averages can be saved either in graphical or numerical formats. Basic channel groups, such as “Left-temporal” and “Right-frontal” are available as default for both MEG and EEG data, but custom channel groups can too be defined.

### Other functionalities

#### Action log and reproducibility

All of the actions used to run analysis steps are part of a logging system. When running an analysis step such as filtering, the user is first asked what parameters to use. When the parameters are selected and the analysis step is run, the name of the action, the selected parameters and a timestamp are recorded to a log. The log is subject-specific and all of the records that have been run within the experiment can be inspected in the past actions dialog (see Fig A1). The dialog shows the records organized by subjects and in temporal order, making it simple to see what has been done before, also for a researcher who has not run the original analysis. This also makes it possible to replicate the analysis of a Meggie experiment elsewhere. It is currently not possible to directly run the analysis steps without the graphical user interface. However, the software is open source, and thus it is also possible to read the implementations of each action, which oftentimes just wrap the mne-python function calls in a multi-subject way.

**Figure A1.**
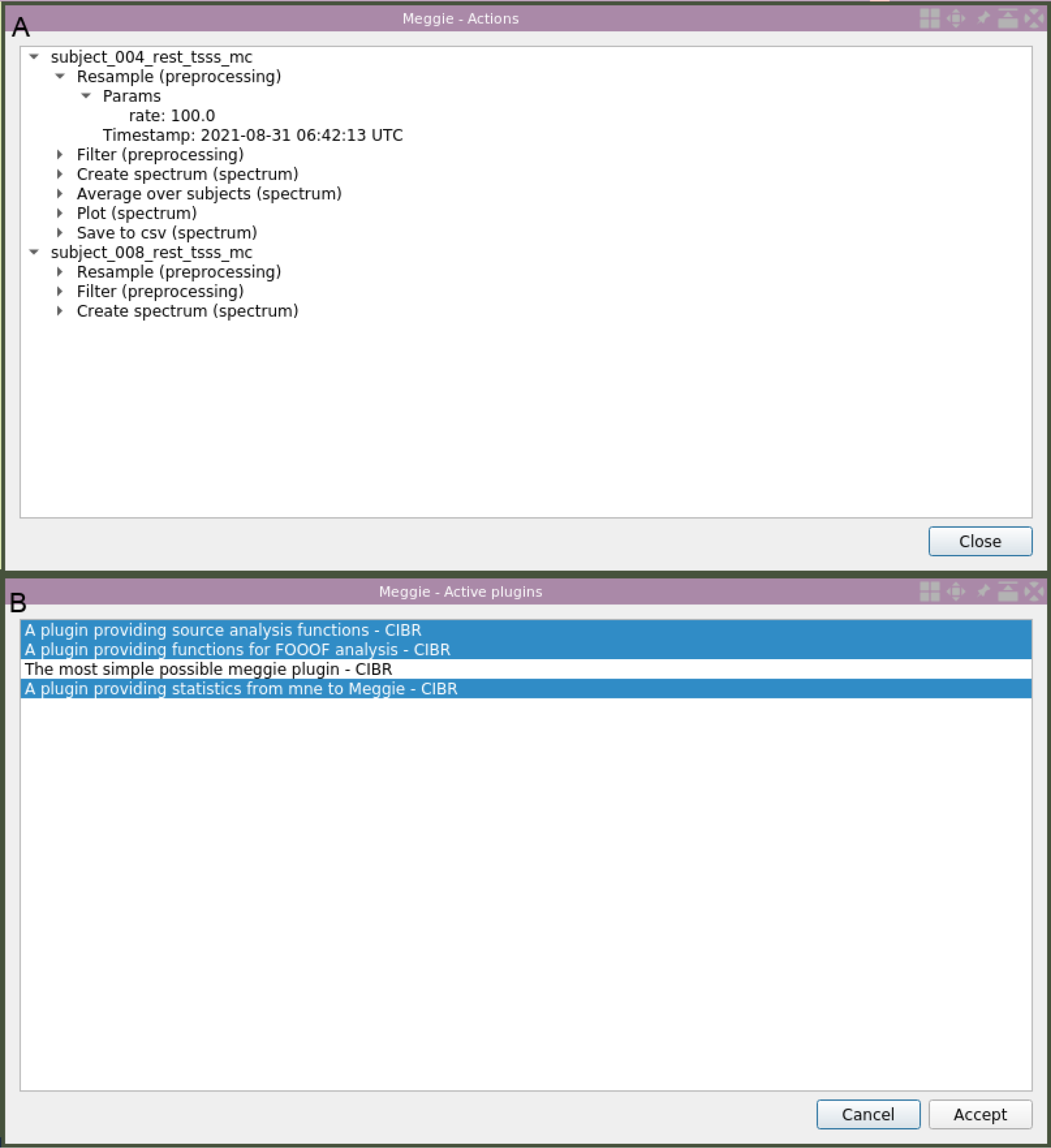
Dialogs for browsing past actions (A) and setting active plugins (B).

#### Multi-subject support

The good support for multiple subjects is perhaps the most important of Meggie. This is realized in the extensive batching capabilities and the methods for combining the data across subjects. Most of the analysis steps (excluding ICA) can be run simultaneously for all subjects of one experiment. In practice this is done by specifying the parameters of an action for a single subject (e.g. what event codes should the epoch collection be based on) and then selecting the action to be applied to every subject with the same settings. The results of all pipelines, i.e. evoked responses, TFRs and spectrums, can be combined across the subjects with grand averages.

## Conclusions

The project was first started because there was a need for more versatile analysis approaches for neuroscience specialists without a programming background. Using a software which provides standard pipelines, provides good default values for parameters, has a good multi-subject support, and stores the used analysis steps with the parameters in one place for reporting, is efficient and fast. In addition to enabling analysis for people without background in programming, it enables analysis for people with background in programming but a limited background in neuroscience. When constructed with care, the GUI may guide the researcher to apply analysis steps in correct order with reasonable default parameters.

The project is actively maintained in CIBR, Jyväskylä, Finland. Yet, we are happy to accept any contributions. The project has been set up in Github, and instructions for participation can be found there. If experiencing any trouble or having ideas for future development, please set up an issue and we can figure out together what is the best way to proceed. Another way to participate is by creating and experimenting with plugins. Instructions for creating and using plugins can be found from the website. It can be a nice way to share your custom methodologies for a wider audience.

Some plugins are already available for use. The statistical analysis plugin, meggie_statistics, provides permutation cluster tests for each of the three basic pipelines. In short, permutation cluster tests are a way to correct for multiple comparisons when running a massive amount of correlated univariate tests with a known correlation structure (Maris and Oostenveld, 2007). The implemented tests are run with subjects as samples and within- and between-subjects designs are supported. At the time of writing, the plugin for inverse modeling, meggie_sourceanalysis, is under development but available for test use. The plan is to implement the three main pipelines, which can currently only work with sensor-level data, to source-level data too.

Meggie answers the demand for easy-to-use and extendable python-based graphical user interface that provides an end-to-end analysis environment for M/EEG data analysis. It is freely available at https://github.com/cibr-jyu/meggie under the BSD license. Installation instructions, documentation and tutorials are found on that website.

## Acknowledgements

This work was supported by the Doctoral School of the Faculty of Information Technology of the University of Jyväskylä. We thank all the great personnel of CIBR who have been involved in the creation, testing and use of the software.

## Supplementary materials

### Architecture and plugins

The way Meggie is programmatically built is very simple. The core consists of Experiment class, which implements logic for saving and loading experiments. It also stores subjects within it, who are the responsibility of a Subject class. The Subject class implements logic for saving and loading subjects, most importantly the corresponding datasets. Subjects can also store different kinds of objects based on predefined datatypes, for example epochs or spectrums. These objects also implement their own logic for saving and loading. The graphical user interface is implemented with Qt5. The most important piece is the main window, which contains the left panel, the bottom console, the menus at the top, and the container for tabs (See Fig 2).

The actual contents of Meggie are implemented as internal plugins, which are functionally no different from external plugins. A plugin is a python package that can introduce datatypes, pipelines and actions (see Fig A1B for plugin selection dialog). Datatypes are classes whose instances can be stored within subjects. Meggie comes natively with four datatypes: Epochs, Evoked, Spectrum and TFR. Actions implement analysis steps by declaring a python code that is run when a button in the right panel of the main view is pressed. A typical button click results in opening a dialog for collecting relevant parameters. When the parameters are accepted (and, if batching, the subjects for which the action is applied are selected), the actual action (where something is transformed or created) begins. Pipelines consist of a declaration of a set of actions organized in tabs. In addition to buttons (actions), tabs may also contain elements for inputs, outputs or additional information. From the declarations meggie dynamically creates the contents of the right main view panel, figures out its layout and fills in the contents. More information about the development in Meggie can be found in the software website https://github.com/cibr-jyu/meggie.

## References

Delorme, A., Makeig, S., 2004. EEGLAB: an open source toolbox for analysis of single-trial EEG dynamics including independent component analysis. J. Neurosci. Methods 134, 9–21. https://doi.org/10.1016/j.jneumeth.2003.10.009

Dressler, O., Schneider, G., Stockmanns, G., Kochs, E.F., 2004. Awareness and the EEG power spectrum: analysis of frequencies. BJA Br. J. Anaesth. 93, 806–809. https://doi.org/10.1093/bja/aeh270

Gramfort, A., Luessi, M., Larson, E., Engemann, D.A., Strohmeier, D., Brodbeck, C., Parkkonen, L., Hämäläinen, M.S., 2014. MNE software for processing MEG and EEG data. NeuroImage 86, 446–460. https://doi.org/10.1016/j.neuroimage.2013.10.027

Gross, J., Baillet, S., Barnes, G.R., Henson, R.N., Hillebrand, A., Jensen, O., Jerbi, K., Litvak, V., Maess, B., Oostenveld, R., Parkkonen, L., Taylor, J.R., van Wassenhove, V., Wibral, M., Schoffelen, J.-M., 2013. Good practice for conducting and reporting MEG research. Neuroimage 65, 349–363. https://doi.org/10.1016/j.neuroimage.2012.10.001

Hari, R., Salmelin, R., 2012. Magnetoencephalography: From SQUIDs to neuroscience: Neuroimage 20th Anniversary Special Edition. NeuroImage, NEUROIMAGING: THEN, NOW AND THE FUTURE 61, 386–396. https://doi.org/10.1016/j.neuroimage.2011.11.074

Hohaia, W., Saurels, B.W., Johnston, A., Yarrow, K., Arnold, D.H., 2022. Occipital alpha-band brain waves when the eyes are closed are shaped by ongoing visual processes. Sci. Rep. 12, 1194. https://doi.org/10.1038/s41598-022-05289-6

Li, Y., Lou, B., Gao, X., Sajda, P., 2013. Post-stimulus endogenous and exogenous oscillations are differentially modulated by task difficulty. Front. Hum. Neurosci. 7.

Maris, E., Oostenveld, R., 2007. Nonparametric statistical testing of EEG- and MEG-data. J. Neurosci. Methods 164, 177–190. https://doi.org/10.1016/j.jneumeth.2007.03.024

Morales, S., Bowers, M.E., 2022. Time-frequency analysis methods and their application in developmental EEG data. Dev. Cogn. Neurosci. 54, 101067. https://doi.org/10.1016/j.dcn.2022.101067

Morgan-Short, K., Tanner, D., 2014. Event-related potentials (ERPs), in: Research Methods in Second Language Psycholinguistics. pp. 127–152. https://doi.org/10.4324/9780203123430

Oostenveld, R., Fries, P., Maris, E., Schoffelen, J.-M., 2010. FieldTrip: Open Source Software for Advanced Analysis of MEG, EEG, and Invasive Electrophysiological Data. Comput. Intell. Neurosci. 2011, e156869. https://doi.org/10.1155/2011/156869

Tadel, F., Baillet, S., Mosher, J.C., Pantazis, D., Leahy, R.M., 2011. Brainstorm: A User-Friendly Application for MEG/EEG Analysis. Comput. Intell. Neurosci. 2011, e879716. https://doi.org/10.1155/2011/879716

Thigpen, N.N., Keil, A., 2017. Event-Related Potentials, in: Reference Module in Neuroscience and Biobehavioral Psychology. Elsevier. https://doi.org/10.1016/B978-0-12-809324-5.02456-1

Vanni, S., Revonsuo, A., Hari, R., 1997. Modulation of the Parieto-Occipital Alpha Rhythm during Object Detection. J. Neurosci. 17, 7141–7147. https://doi.org/10.1523/JNEUROSCI.17-18-07141.1997

Zhao, L., He, Y., 2013. Power Spectrum Estimation of the Welch Method Based on Imagery EEG. Appl. Mech. Mater. 278–280, 1260–1264. https://doi.org/10.4028/www.scientific.net/AMM.278-280.1260

Zhigalov, A., Heinilä, E., Parviainen, T., Parkkonen, L., Hyvärinen, A., 2019. Decoding attentional states for neurofeedback: Mindfulness vs. wandering thoughts. NeuroImage 185, 565–574. https://doi.org/10.1016/j.neuroimage.2018.10.014

